# Spindle scaling promotes adaptation to polyploidy

**DOI:** 10.64898/2026.08.17.745319

**Authors:** Gabriel Cavin, Cordell Clark, Liliana Piñeros, Donggeon Woo, Taejoon Kwon, Rebecca Heald

## Abstract

How the cell division machinery adapts to increases in genome copy number (ploidy) remains an open question with implications for evolution and cancer. Using *Xenopus* egg extracts and species spanning 2N to 12N ploidy, we found that stepwise increases in ploidy drove spindle multipolarity, accompanied by larger spindles, increased microtubule density, and differential scaling of spindle assembly factor localization. In contrast, extracts from the dodecaploid species *Xenopus longipes* supported robust bipolar spindle assembly at all ploidies. Engineering *X. laevis* extracts to mimic *X. longipes* by modulating microtubule regulators frequently overexpressed in cancer, or confining spindles inside small droplets, decreased spindle length and significantly rescued bipolarity. Thus, tuning spindle microtubule dynamics and architecture enables adaptation to increased chromosome number across acute and evolutionary timescales.

## Introduction

Polyploidy, the presence of more than two complete sets of chromosomes, poses a fundamental challenge to faithful cell division because spindle microtubules must properly attach to and segregate at least twice as many chromosomes. Although polyploidy is widespread across plants, amphibians, and fish, stable whole-organism polyploidy is absent in extant mammals^1,2,3^. Nevertheless, polyploid cells are common in mammalian somatic tissues. While some polyploid cell types are terminally differentiated, others, including hepatocytes, retain proliferative capacity^4^ but their divisions are frequently associated with transient multipolar spindle formation that leads to chromosome segregation errors and aneuploidy^5,6^, highlighting the physiological constraints imposed by increased ploidy in vertebrate systems.

Polyploidy is also common in cancer, where whole-genome doubling (WGD) occurs early during the evolution of many carcinomas^7,8,9,10^. Tetraploid cells arising from cytokinesis failure or mitotic slippage possess supernumerary centrosomes, the microtubule-organizing centers that form spindle poles in most animal cells. Extra centrosomes promote transient multipolar spindle formation, erroneous microtubule attachments to chromosomes at kinetochores, and aneuploidy. While this instability fuels tumor evolution by generating genetic diversity, excessive chromosomal instability can trigger cell death, indicating that polyploid cancer cells must acquire adaptations to remain viable. One mechanism by which cells avoid lethal multipolar divisions is to cluster extra centrosomes or supernumerary spindle poles to restore bipolar spindle formation^11,12,13,14,15^. However, even when centrosomes are initially clustered, excess chromatin can form a physical barrier that impedes pole coalescence and bipolar spindle assembly^16^. Furthermore, tetraploidy drives chromosomal instability in mouse embryos, where early embryonic divisions normally occur in the absence of centrosomes.^17^ Consistent with the need to manage these spindle assembly challenges, cancer genome analyses and functional studies have identified mitotic vulnerabilities in WGD cells, including a dependence on the kinesin-8 motor KIF18A, which suppresses kinetochore microtubule dynamics to promote accurate chromosome alignment and segregation^18^.

More broadly, the repeated emergence of polyploid lineages across the tree of life suggests that cells can adapt to increased genome content, yet the mechanisms that enable faithful chromosome segregation in polyploids remain poorly understood^19^. Experiments in budding yeast have revealed intrinsic constraints on mitotic spindle architecture, as the number of spindle microtubules does not scale with increasing ploidy, promoting erroneous kinetochore– microtubule attachments^20,21^. The limits of vertebrate mitotic spindle scaling are less clear. Cell size scales with genome size, and spindle and nuclear size scale with cell size, potentially preserving overall geometry^22,23^. However, how the spindle itself evolves to accommodate increased chromosome number is unknown. Most vertebrate studies have relied on cancer-derived or experimentally induced polyploid cells, making it difficult to distinguish the immediate consequences of increased chromosome number from adaptations that enable cells to accommodate the polyploid state. Defining these adaptations will reveal fundamental principles of spindle function and may uncover therapeutic vulnerabilities in polyploid cancers.

Here we take advantage of the *Xenopus* genus, which spans a broad range of naturally occurring ploidies from diploid *X. tropicalis* to dodecaploid *X. longipes*—a small, highly endangered species that evolved in a crater lake in Cameroon, Africa^24^. The capacity of cytoplasmic extracts prepared from unfertilized *Xenopus* eggs to support robust, centrosome-independent spindle assembly *in vitro* provides a powerful system for dissecting intrinsic spindle architectural adaptations to increased genome copy number^25,26,27^. By combining nuclei and cytoplasm from different species, we distinguish the immediate effects of increased ploidy from evolved cytoplasmic adaptations and, through comparative and functional analysis of spindle assembly across species, identify biochemical and physical mechanisms that enable the cell division machinery to accommodate multiple genome copies and mitigate a key barrier to polyploidization.

## Results

### Increased ploidy lowers the efficiency of bipolar spindle assembly

To manipulate chromosome number during spindle assembly, we used the *Xenopus* egg extract system (Figure 1A), in which cytoplasm extracted from metaphase II-arrested diploid *X. tropicalis* (2N) or allotetraploid *X. laevis* (4N) eggs is combined with a DNA source to initiate spindle formation^28^. We isolated nuclei spanning multiple ploidies from *Xenopus* embryos at Nieuwkoop and Faber (NF) stage 8 (4,000–8,000 cells), confirmed chromosome content by metaphase spreads (Supplemental Figure 1), and assembled spindles around nuclei of matched or mismatched ploidy in *X. tropicalis* or *X. laevis* extracts (Figure 1B). In addition to 2N, 4N, and 12N ploidies, 3N and 6N embryonic nuclei were generated by cold-shocking zygotes to prevent second polar body extrusion. With increasing chromosome number, overall acentrosomal meiotic spindle morphology was maintained, but spindle size increased and spindle bipolarity declined significantly with each increase in genome copy number, falling from 83% in species-matched controls to 29% and 39% in the highest-ploidy conditions in *X. tropicalis* and *X. laevis* extracts, respectively (Figure 1C). In contrast, reducing ploidy or changing the concentration of nuclei did not affect spindle bipolarity (Supplemental Figure 2), indicating that the defects are specific to chromosome gain and arise autonomously during spindle assembly.

**Fig 1.**
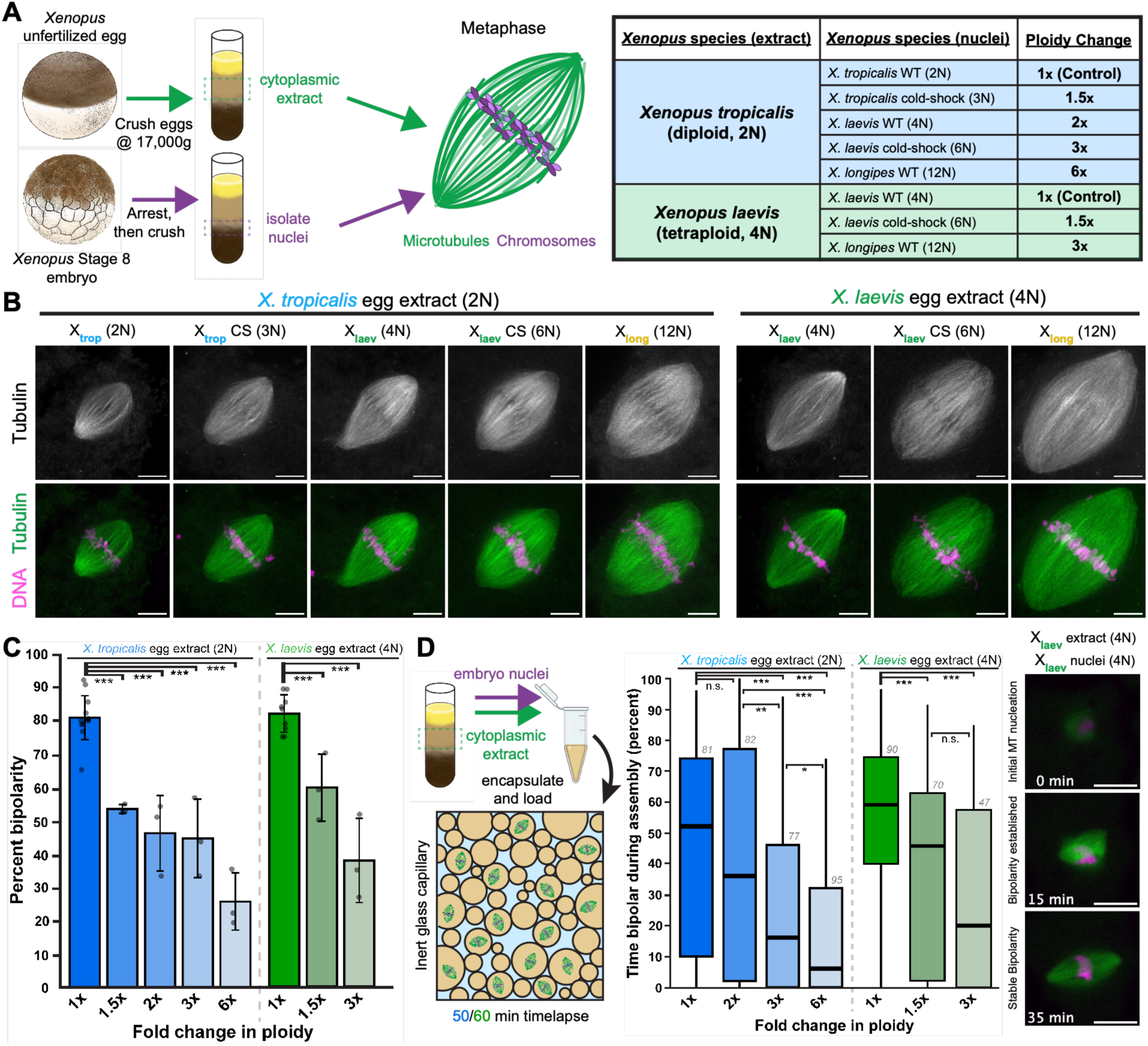
Increasing ploidy increases spindle size and reduces bipolarity. **(A)** Workflow for assembling spindles in *Xenopus* egg extracts using nuclei of varying ploidy. **(B)** Fixed spindle squashes showing all extract-nuclei combinations tested, CS denotes cold-shock of zygote that increases ploidy by 50% by blocking polar body extrusion. Microtubules are shown in green and DNA in magenta. Scale bar, 10 µm, all images are maximum-intensity projections of z-stacks. **(C)** Quantification of spindle bipolarity for the extract-nuclei combinations shown in (B). Each point represents one experimental replicate (≥20 spindles per replicate), bars indicate the mean ± standard deviation. Asterisks denote p values determined by ANOVA. **(D)** Workflow for high-throughput live imaging of spindle assembly. For each spindle, the fraction of time spent in a bipolar state was measured during 50 or 60 minutes for *X. laevis* and *X. tropicalis* extract, respectively. Numbers above boxplots indicate total number of spindles analyzed per condition. Scale bar, 25 µm. All images are single optical sections. Spindle counts for all conditions are provided in Table S1.Figure 2

To determine whether reduced bipolarity reflected increased ploidy rather than species incompatibilities^29^, we generated additional spindle-nuclei combinations (Supplemental Figure 3). Haploid *X. laevis* embryos generated with UV-irradiated sperm produced 2N nuclei that assembled bipolar spindles in *X. tropicalis* extracts at frequencies comparable to 2N *X. tropicalis* nuclei. Likewise, 4N nuclei isolated from allotetraploid *Xenopus borealis* embryos assembled bipolar spindles in *X. laevis* extract at frequencies similar to 4N *X. laevis* nuclei. In addition, *X. borealis* egg extracts^30^ yielded spindle bipolarity levels comparable to *X. laevis* extracts across the same ploidy series. Together, these controls indicate that reduced spindle bipolarity is driven by increased chromosome number.

Live imaging further revealed that spindles remained highly dynamic and did not become trapped in persistent multipolar states; instead, increasing ploidy progressively destabilized spindle bipolarity, with the average fraction of time spent bipolar falling from 54% and 59% in species-matched controls to 6% and 20% at the highest ploidy in *X. tropicalis* and *X. laevis* extracts, respectively (Figure 1D). Similar dynamic transitions through multipolar states have been observed in newly formed tetraploid cells^13,16^.

### Abrupt increases in ploidy alter spindle architecture and expand the Ran-GTP gradient

Although the percentage of bipolar spindles formed declined similarly in *X. tropicalis* and *X. laevis* extracts at equivalent fold increases in ploidy, bipolar spindle architecture responded differently in the two species. Morphometric analysis revealed that spindle length and width increased with ploidy in both extracts (Figure 2A; Supplemental Figure 4). Notably, spindle width exhibited strong scaling with ploidy, in agreement with previous observations of chromosome-dependent spindle-width scaling in *Xenopus* eggs, *C. elegans* embryos, and across eukaryotes^24,31,32^. However, whereas spindle shape was largely maintained in *X. tropicalis* extracts, *X. laevis* extract spindles became increasingly wide relative to their length. In addition, spindle poles became progressively less focused with increasing ploidy in *X. laevis* extracts, while *X. tropicalis* spindles retained focused poles except at the highest ploidy.

**Fig 2.**
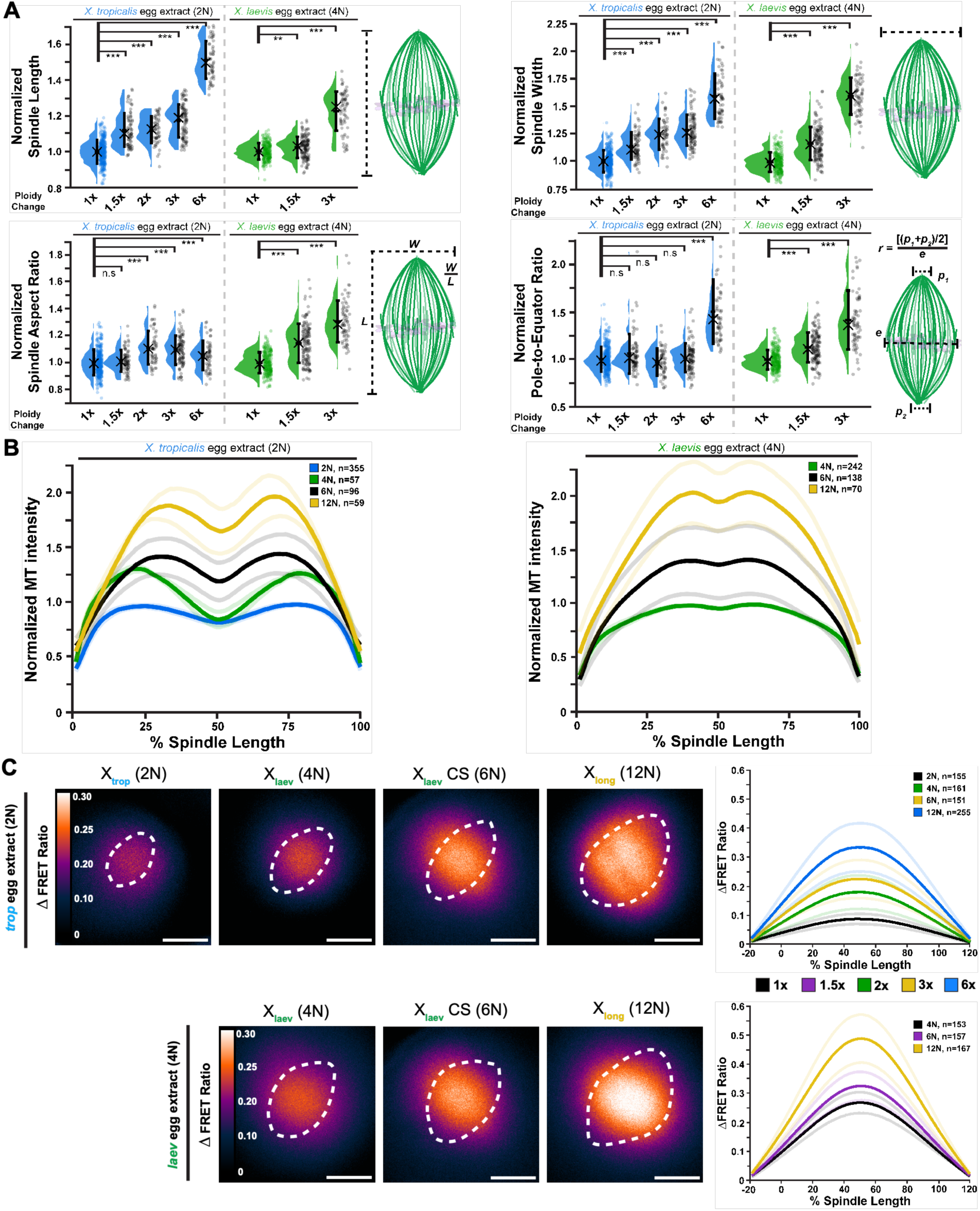
Increasing ploidy alters spindle morphology and expands the Ran-GTP gradient. **(A)** Measurements of bipolar spindle morphological features including spindle length, width, aspect ratio, and ratio of the spindle pole to equator width as a measure of pole focusing. Each point represents a single spindle. Data are pooled from at least three biological replicates (≥55 spindles per replicate). Bars indicate the mean ± standard deviation and crosses indicate the median. Asterisks denote p values determined by ANOVA; n.s., not significant. Spindle numbers for each measurement are provided in Table S1. For additional spindle measurements, see Supplementary Figure 4. **(B)** Linescan analysis of microtubule intensity along the pole-to-pole spindle axis. Each solid line represents the mean profile generated from individual linescans normalized to spindle length. Lighter lines of same color represent ± SEM. **(C)** Live FRET imaging of spindles in *X. tropicalis* and *X. laevis* egg extracts using the Rango biosensor to visualize the Ran-GTP gradient surrounding chromosomes. The dashed line denotes the spindle boundary (imaged in separate channel). The calibration bar indicates changes in the FRET/CFP ratio. Each solid line represents the mean Ran-GTP profile generated from individual linescans, normalized to percent spindle length. Lighter lines of same color represent ± SEM. Scale bar, 25 µm. In (B) and (C), the number of spindles analyzed per ploidy is shown.

In both extracts, higher ploidy increased spindle tubulin fluorescence, indicating greater microtubule density (Figure 2B). Because chromosome-bound RCC1 generates the Ran-GTP gradient that promotes microtubule nucleation^33^, we reasoned that additional chromosomes would enhance local Ran signaling. Consistent with this prediction, the Ran-GTP gradient broadened and intensified with ploidy, with both parameters scaling approximately linearly with chromosome number (Figure 2C). Together, these results suggest that increased ploidy enhances Ran-dependent microtubule assembly, altering spindle architecture and destabilizing bipolarity. Species-specific responses may reflect differences in spindle assembly mechanisms, as *X. tropicalis* spindles are less dependent on Ran signaling, contain higher levels of TPX2, and exhibit greater microtubule-destabilizing activity than *X. laevis* spindles^34,35^.

### Dodecaploid *Xenopus longipes* uncouples spindle scaling from chromosome number

The spindle bipolarity defects caused by acute increases in ploidy would be unlikely to support the viability and propagation of a stable polyploid lineage *in vivo*, yet the existence of the dodecaploid (12N) *X. longipes* demonstrates that mitotic adaptation to polyploidy can occur^24^. Thus, we examined native *X. longipes* spindle assembly to identify features that may enable the spindle to accommodate a highly polyploid genome. *X. longipes* egg extracts assembled robust bipolar spindles with acentrosomal morphology and assembly kinetics similar to the other *Xenopus* species, sustaining high bipolarity across ploidy levels, including their native dodecaploid genome content (Figure 3; Supplemental Figure 5). Despite having a shorter pole-to-pole length than native *X. laevis* extract spindles, *X. longipes* spindles exhibited the greatest spindle width, metaphase plate width, and spindle volume among the three *Xenopus* species (Supplemental Figure 6). However, compared with mixed-ploidy spindles assembled around *X. longipes* nuclei in *X. tropicalis* and *X. laevis* extracts, spindles formed from dodecaploid nuclei in native *X. longipes* extract were significantly smaller and possessed more tightly focused poles (Figure 3C). Although a Ran-GTP gradient was present around *X. longipes* chromosomes in native extract, its strength was intermediate between native extract-nuclei combinations of the other *Xenopus* species and its spatial profile was flatter than that of *X. laevis* despite having threefold more chromosomes (Figure 3D). Thus, whereas spindle size and Ran-GTP gradient amplitude scale with ploidy in *X. laevis* and *X. tropicalis* egg extracts, both are attenuated relative to chromosome number in *X. longipes*. These findings suggest that *X. longipes* has evolved mechanisms that decouple spindle size scaling from chromosome-dependent Ran signaling.

**Fig 3.**
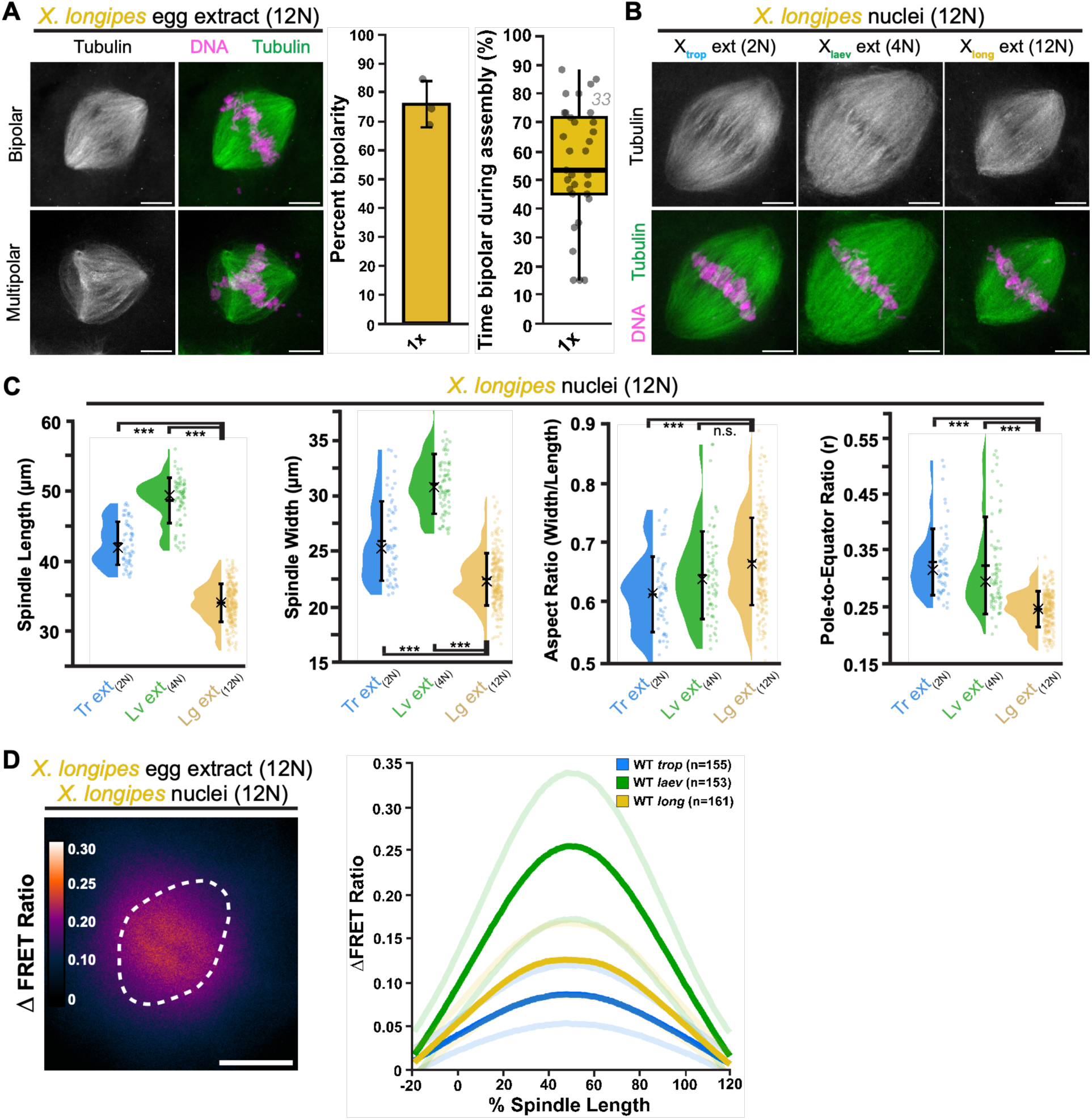
Egg extracts from dodecaploid *Xenopus longipes* support robust bipolar spindle assembly around polyploid nuclei. **(A)** Spindle bipolarity of native *X. longipes* egg extract spindles. Fixed spindle squashes show microtubules in green, DNA in magenta. For percent bipolarity of fixed spindles, each point represents one experimental replicate (≥20 spindles per replicate). For the percent time spindles spent in a bipolar state, each point represents a single spindle (33 total) over 60 minutes of a timelapse video. Bars indicate the mean ± standard deviation. **(B)** Fixed squashes of spindles formed with *X. longipes* nuclei (12N) in extracts of all three species. For (A) and (B), scale bar = 10 µm, all images are maximum-intensity projections of z-stacks. **(C)** Spindle morphometrics from spindles observed in (B) with *X. tropicalis* and *X. laevis* data adapted from Figure 2A. Each point represents one spindle with data pooled from at least three biological replicates (≥30 spindles per replicate). Bars indicate mean value ± standard deviation and crosses indicate the median. Asterisks denote p values determined via ANOVA. **(D)** Live FRET imaging of native *X. longipes* spindle with Rango biosensor to visualize the Ran-GTP gradient surrounding chromosomes. The dashed line denotes the spindle boundary (imaged in separate channel). The calibration bar indicates changes in the FRET/CFP ratio. Linescan analysis shows the relative profile of the Ran-GTP gradient in native (WT) spindles assembled in extracts of all three *Xenopus* species with *X. tropicalis* and *X. laevis* data adapted from Figure 2C. Each solid line represents the mean Ran-GTP profile generated from individual linescans, normalized to percent spindle length. Lighter lines of same color represent ± SEM. Scale bar, 25 µm. The number of spindles analyzed per species is shown.

### KIF2A recruitment is enhanced in *Xenopus longipes*

Our observations of this decoupling motivated us to further dissect how acute versus evolutionary increases in ploidy affect spindle composition. We used immunofluorescence to survey spindle-associated proteins in mixed-ploidy reactions and in native dodecaploid *X. longipes* extracts (Figure 4; Supplemental Figures 7-10). The proteins examined have diverse roles in spindle assembly and function: the kinesin-5 Eg5 and kinesin-14 XCTK2 (HSET) motors sort microtubules into a bipolar array and focus spindle poles, respectively, while chromatin-bound RCC1 generates the Ran-GTP gradient that promotes microtubule nucleation. Additional proteins examined included the kinetochore-associated kinesin-7 CENP-E, which contributes to chromosome alignment, and three factors previously implicated in spindle scaling and microtubule dynamics: the kinesin-13 KIF2A, the microtubule-severing enzyme Katanin, and the multifunctional spindle assembly factor TPX2 ^34,35,36^. In both *X. tropicalis* and *X. laevis* extracts, Eg5, XCTK2, CENP-E, Katanin, TPX2, and RCC1 scaled proportionally with ploidy and spindle microtubule density as measured by tubulin intensity (Figure 4A; Supplemental Figures 7-9), whereas KIF2A did not. Spindle KIF2A levels remained relatively constant as ploidy and spindle size increased, resulting in a progressive decrease in the KIF2A:tubulin ratio (Figure 4B). Thus, unlike the other spindle assembly factors examined, KIF2A recruitment failed to scale with the increased microtubule mass generated by increasing ploidy.

**Fig 4.**
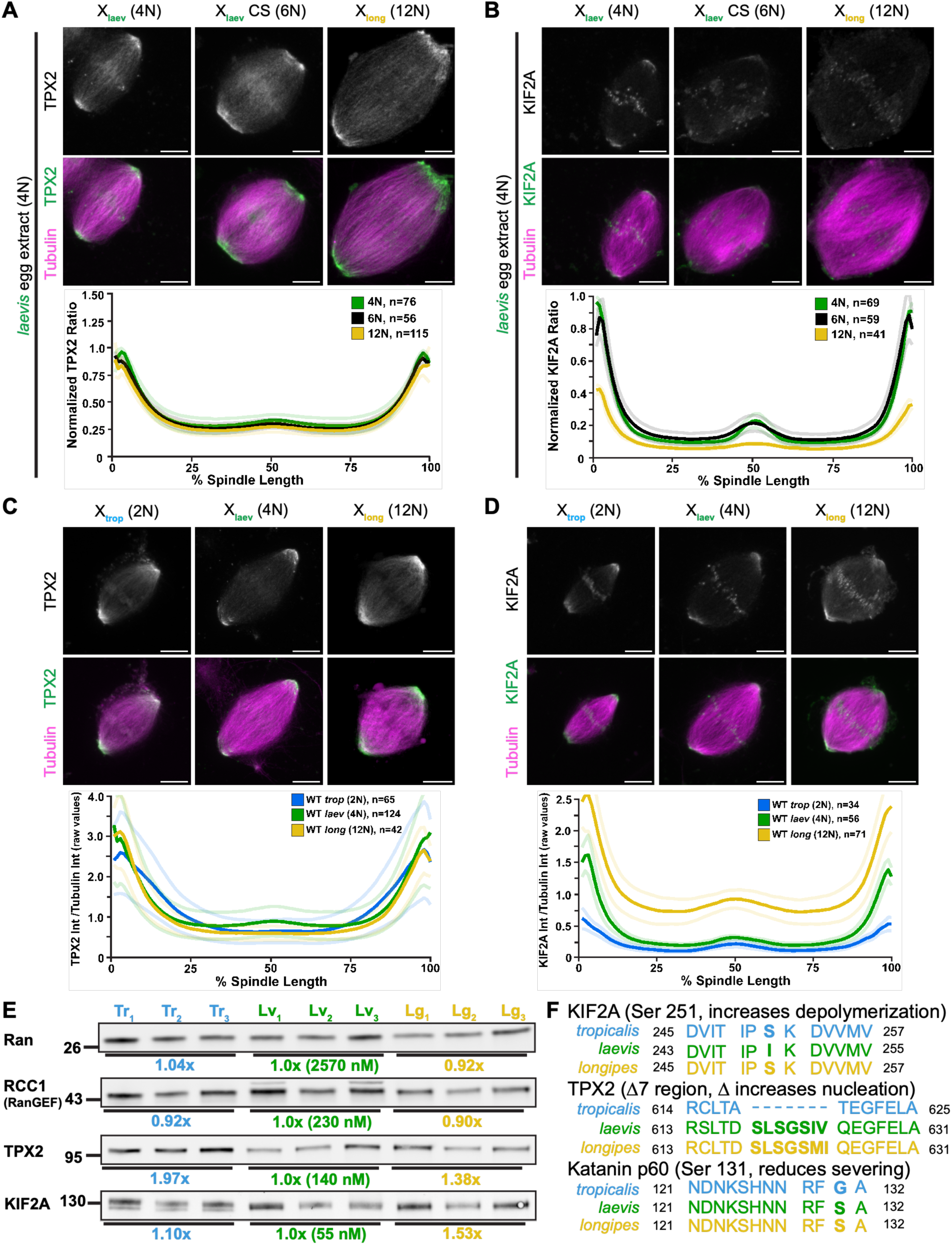
Increasing ploidy differentially affects TPX2 and KIF2A localization. **(A,B)** Linescan analysis of TPX2 and KIF2A intensity relative to spindle microtubule intensity along the pole-to-pole axis in *X. laevis* extracts (normalized to the species-matched reaction). Solid lines represent the average profile, and shaded lines of same color represent ± SEM. The number of individual linescans in each condition is shown. For the same experimental conditions in *X. tropicalis* extracts, see Supplemental Figure 7. Representative images for all panels are sum projections of z-stacks. Scale bar, 10 µm. **(C,D)** Linescan analysis of TPX2 and KIF2A intensity relative to spindle microtubule intensity along the pole-to-pole axis of native WT *Xenopus* spindles of the three species. Solid lines represent the average profile generated from individual linescans (n shown). Shaded lines represent ± SEM. All images are sum projections of z-stacks. Scale bar = 10 µm. **(E)** Western blotting of various proteins in all three *Xenopus* egg extracts. All lanes were loaded with 10 µg total protein. The average intensity of *X. laevis* bands was used to normalize the other species, with estimated concentrations in parentheses. **(F)** Multiple sequence alignment of three proteins (TPX2, KIF2A, and Katanin p60) across all three *Xenopus* species. *X. longipes* sequences were obtained by *de novo* transcriptome assembly of total mRNA isolated from organs of an adult *X. longipes* female.

We therefore asked whether KIF2A recruitment is altered in native *X. longipes* cytoplasm. The localization of spindle-associated proteins in native *X. longipes* spindles resembled that observed in the other *Xenopus* species (Supplemental Figure 10). Comparison of spindles assembled in extracts from each species at their native ploidy revealed significantly more KIF2A relative to tubulin in *X. longipes* than in either *X. tropicalis* or *X. laevis*, whereas other spindle assembly factors, including TPX2, were present at comparable relative levels (Figure 4C-D). Together, these findings identify KIF2A as a factor whose recruitment is uncoupled from microtubule density following acute increases in ploidy but selectively enhanced in *X. longipes*, suggesting that increased microtubule depolymerization at spindle poles may contribute to shorter spindle length and adaptation to increased ploidy.

Consistent with enhanced KIF2A recruitment, Western blot analysis showed that although the levels of most spindle proteins examined were similar across species (Supplemental Figure 11), *X. longipes* eggs contained approximately 50% higher levels of KIF2A, and intermediate levels of TPX2 between *X. laevis* and *X. tropicalis* (Figure 4E). Notably, levels of the kinesin-13 MCAK and levels of KIF18A, a kinesin-8 required for mitotic fidelity in WGD cancer cells^18^, were not increased in *X. longipes* compared with *X. laevis* (Supplemental Figure 11). Protein activity can also be altered through changes in primary sequence or post-translational regulation; sequence alignment revealed that many *X. longipes* spindle proteins were nearly identical to *X. laevis* homologs. However, *X. longipes* KIF2A retains an activating phosphorylation site absent from *X. laevis*, potentially enhancing microtubule depolymerase activity (Figure 4F). Together, these differences identify KIF2A abundance and spindle recruitment, along with altered TPX2 levels, as candidate adaptations that may contribute to the relatively short spindles and enhanced bipolarity observed in native dodecaploid extracts.

### Reducing spindle size and increasing microtubule turnover promote bipolar spindle assembly around polyploid nuclei

To determine whether the evolutionary differences observed contribute to spindle adaptation, we modified *X. laevis* extracts to mimic *X. longipes* by supplementing TPX2 and KIF2A to native *X. longipes* levels (Figure 5A-B). This decreased spindle size and substantially rescued bipolarity under conditions of increased ploidy, increasing bipolar spindle assembly from 29% to 51% and extending the time spindles spent in a bipolar state from 26% to 56%. Prior studies have shown that increasing microtubule turnover can improve spindle bipolarity and reduce segregation defects in early human embryos and several cancers^18,37,38^. To test whether increased microtubule destabilization more generally promotes bipolarity, we activated the kinesin-13 KIF2C (MCAK) with UMK57^38^. MCAK activation partially rescued spindle bipolarity at increased ploidy in *X. laevis* extracts (Figure 5B-C). However, higher UMK57 concentrations abolished this rescue and reduced bipolarity under species-matched conditions, indicating a requirement for balanced microtubule dynamics. Supplementing TPX2 to native *X. longipes* levels did not enhance rescue at lower UMK57 concentrations, and at high concentrations the combination reduced spindle bipolarity under species-matched conditions. However, TPX2 addition enhanced rescue of spindle bipolarity at a threefold increase in ploidy in *X. laevis* egg extracts (from 28 % to 45%). Nevertheless, no MCAK activation condition restored bipolarity to the same extent as combined KIF2A and TPX2 supplementation. Together, these results support coordinated regulation of TPX2 and KIF2A as a mechanism that promotes robust spindle assembly upon increased ploidy.

**Fig 5.**
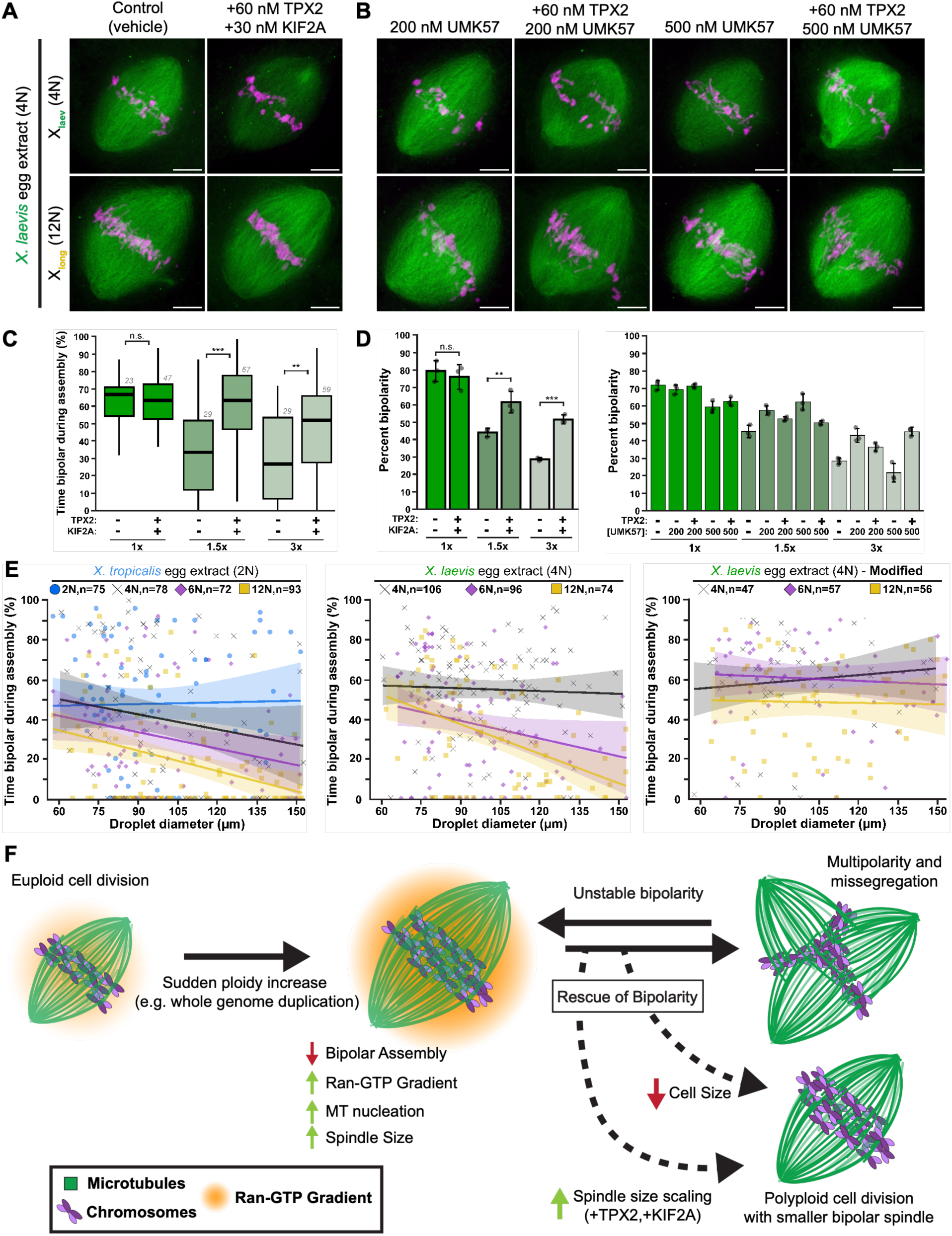
Tuning spindle dynamics and size promotes bipolarity when ploidy is increased. **(A)** Fixed images of *X. laevis* extract spindles with or without purified *X. laevis* TPX2 (60 nM) and KIF2A (30 nM). **(B)** Fixed images of *X. laevis* extract spindles with or without purified *X. laevis* TPX2 (60 nM) and treatment with the MCAK agonist UMK57 (200 nM or 500 nM). All images are maximum projections. Scale bar, 10 µm. **(C)** Quantification of time spent in a bipolar state during live imaging, displayed as boxplots. The number above each boxplot denotes the number of spindles measured in each condition. **(D)** Quantification of the percentage of bipolar spindles observed in (A,B). Each point represents one experimental replicate (≥25 spindles per replicate), bars indicate the mean ± standard deviation. (C, D) Asterisks denote p values determined by ANOVA. **(E)** Relationship between spindle bipolarity during assembly and the diameter of the droplet in which the spindle forms. Each point represents one complete timelapse movie of a droplet. Regression lines are colored to match their respective data points. Each condition contains droplets from at least three biological replicates (≥10 timelapses per replicate). The total number of droplets for each condition is shown. **(F)** Proposed model synthesizing the observations made in this work. Sudden increases in ploidy directly expand and strengthen the Ran-GTP gradient during spindle assembly, driving microtubule nucleation near chromosomes. This surge in microtubule nucleation can be counteracted by tuning spindle dynamics and size, limiting the stabilization of nascent poles and promoting their coalescence into a robust bipolar spindle.

Beyond changes in spindle protein levels and activities, spindle size and architecture are also influenced by cell size, which decreases drastically during the rapid cleavage divisions of early embryogenesis. Previous studies have shown that spindle size scales with cell size, in part because spindle components such as tubulin become limiting as cytoplasmic volume decreases^39,40^. To determine whether cytoplasmic volume influences spindle robustness at increased ploidy, we re-analyzed our live imaging data as a function of droplet diameter (Figure 5D). Spindles at the highest ploidy increases spent significantly more time bipolar in small droplets than in larger droplets (47% in 60-100 µm droplets versus 12% in droplets >100 µm), whereas droplet size had no effect under species-matched conditions. Moreover, modifying *X. laevis* extract to mimic native *X. longipes* conditions by increasing levels of TPX2 and KIF2A eliminated the relationship between droplet size and spindle bipolarity across ploidy conditions. These results identify reduced cytoplasmic volume as a physical mechanism that constrains spindle size and promotes bipolarity at increased ploidy, while evolutionary changes in *X. longipes* render spindle assembly robust to variation in both chromosome number and cytoplasmic volume.

## Discussion

By varying chromosome number while maintaining a constant cytoplasmic environment, we assessed the immediate effects of increasing ploidy on spindle assembly. Increased ploidy in *X. tropicalis* and *X. laevis* egg extracts strengthened the Ran-GTP gradient, increased microtubule density and spindle size, and decreased spindle bipolarity. These findings suggest that multipolar spindles, a major source of chromosome mis-segregation and aneuploidy in cancer, can arise not only from centrosome amplification but also from excess spindle microtubules that cannot be efficiently organized into two poles. In contrast, the dodecaploid *X. longipes* has evolved mechanisms that limit spindle length despite its greatly expanded genome. One adaptation appears to involve a weaker Ran-GTP gradient relative to chromosome number. Because Ran and RCC1 abundance and localization are similar across the three frog species, this difference may reflect altered RCC1 or RanGAP activity, or other changes in Ran regulation. Downstream of Ran, enhanced recruitment of the microtubule-depolymerizing kinesin KIF2A to spindle poles, together with altered TPX2 levels, provides a complementary mechanism to restrain spindle size despite high ploidy. KIF2A likely promotes spindle bipolarity by enhancing microtubule turnover at spindle poles, counterbalancing Ran-dependent microtubule nucleation, limiting spindle length, and facilitating pole focusing. TPX2 is a known spindle-scaling factor^33^ that tunes spindle architecture through its roles in microtubule nucleation and stabilization, Aurora A activation, and Eg5-mediated microtubule sorting^41^. Together, these findings support a model in which adaptation to polyploidy involves rebalancing microtubule assembly and turnover to maintain a spindle architecture compatible with robust bipolarity (Figure 5E).

This mechanism fits a broader principle emerging from studies of spindle scaling. Factors that regulate microtubule dynamics are recurrent developmental and evolutionary regulators of spindle architecture. Katanin and TPX2 contribute to spindle size differences between *X. tropicalis* and *X. laevis*^34,35^, whereas KIF2A is a major spindle-scaling factor in the miniature frog *Hymenochirus boettgeri*^36^ and during early embryonic cleavage divisions^40,42^. Our findings extend this principle to evolutionary adaptation to genome expansion, suggesting that enhanced microtubule depolymerization offsets increased chromosome-driven microtubule nucleation to limit spindle growth and promote bipolarity. Consistent with this model, spindle bipolarity at increased ploidy was improved by KIF2A supplementation or MCAK activation. Independently, reducing cytoplasmic volume also promoted bipolarity by constraining spindle growth, indicating that spindle architecture can be tuned through both biochemical and physical mechanisms. Although both KIF2A and MCAK are kinesin-13 microtubule depolymerases, they have distinct functions and localizations, with KIF2A enriched at spindle poles, whereas MCAK is more broadly distributed and regulates microtubule dynamics throughout both interphase and mitosis^43^. The extent of rescue by KIF2A and TPX2 depended on ploidy, restoring spindle bipolarity nearly completely at 1.5-fold ploidy but only partially at threefold ploidy, suggesting that additional factors contribute to adaptation as chromosome number increases. Although the spindle kinesin-8 KIF18A is required for the viability of WGD cancer cells^18^ and influences spindle length and pole integrity in *X. laevis* egg extracts^44^, its levels were not increased in *X. longipes*, arguing against altered KIF18A abundance as an adaptive mechanism. Together, our results suggest that rather than simply scaling spindle size with genome size, evolutionary remodeling of microtubule dynamics and spindle architecture enables cells to accommodate increased chromosome number while maintaining robust bipolar spindle assembly.

These findings also have implications for whole-genome-doubled (WGD) cancers. WGD cells face two major mitotic challenges: accommodating the greater spindle microtubule mass that can accompany a larger chromosome complement and faithfully segregating those chromosomes. A shorter, more dynamic spindle may help meet these challenges by promoting pole focusing and centrosome clustering while maintaining the microtubule turnover required for chromosome capture and alignment. Consistent with this idea, TPX2 and KIF2A are frequently overexpressed in human cancers^41,45^, increased TPX2 expression correlates with reduced spindle size^46^, and TPX2 is required for the survival of many WGD tumor cells^47^. Strikingly, decreased cell and nuclear size in tetraploid cells is associated with improved mitotic fidelity, increased fitness and tumorigenicity, and poorer clinical outcomes in WGD-positive tumors^48,49^. Our finding that reducing cytoplasmic volume enhanced stable spindle bipolarity provides a potential mechanistic link between reduced cell size and improved mitotic fidelity in tetraploid cancer cells. Because smaller cells contain reduced pools of tubulin and other cytoplasmic components, these factors can become limiting and constrain spindle size, as previously demonstrated by confinement of *Xenopus* egg extracts in small droplets^23^. Thus, reduced cell size may limit the excess microtubule assembly driven by increased chromosome number, promoting a spindle architecture that favors bipolarity. More broadly, spindle-scaling mechanisms that enable adaptation to larger genomes may also provide routes by which WGD cancer cells evolve to maintain the mitotic fidelity necessary for continued proliferation.

## Materials and methods

### Isolation of Stage 8 embryo nuclei and ploidy verification

Embryo nuclei were isolated from NF Stage 8 obtained by in vitro fertilization (IVF) or natural mating. Embryos were arrested in interphase by transfer to 0.1x MMR containing 300 µg/mL cycloheximide at approximately 4.5 h post-fertilization (hpf) *X. tropicalis* (23°C), 6.0 hpf for *X. laevis* (23°C), and 8.0 hpf for *X. longipes* (17°C). After 60 min, embryos were exchanged into Egg Lysis Buffer (ELB) containing 10 µg/mL LPC and 20 µg/mL cytochalasin B, then fractionated in 1.5 mL microcentrifuge tubes as described for *Xenopus* egg extracts^40^. Cytoplasmic and membrane fractions were collected and gently mixed, and nuclei concentration was estimated by counting in a Hoechst-stained 0.5 µL drop. Glycerol was added to 8%, and aliquots were flash-frozen in liquid nitrogen and stored at −80°C.

Triploid embryos were induced by cold shock as described previously^50^. Fertilized embryos were transferred to ice-cold 0.1x MMR at 10 min (*X. tropicalis*) or 13 min (*X. laevis*) post-fertilization for 5 min to disrupt the meiosis II spindle and prevent polar body extrusion (∼80% efficiency), then returned to room temperature 0.1x MMR and maintained according to standard IVF protocols. Haploid embryos were generated by UV irradiation of sperm prior to fertilization^51^. Fresh testes were dissected into L-15 medium containing 10% FBS in a 1.5 mL microfuge tube, homogenized using scissors and a pestle, and centrifuged briefly to remove tissue debris. The sperm-containing supernatant was transferred to a 6 cm glass dish and irradiated in a Stratalinker with one 50,000 J pulse (*X. tropicalis*) or two 30,000 J pulses (*X. laevis*). Irradiated sperm were immediately used for fertilization, and embryos maintained according to standard IVF protocols.

Embryo ploidy was verified by metaphase chromosome spreads from ∼15 sibling embryos collected at NF Stages 36-40 (tailbud). Following anesthesia in 0.1% benzocaine, dorsal tissue was isolated and incubated in 1.2 mg/mL colchicine for 60 min, distilled water for 20 min, and 70% acetic acid for 5 min. Tissue was transferred to glass slides, excess liquid was removed with filter paper, and then covered with a coverslip. Slides were compressed under a 10 kg weight for 5 min, the weight lifted directly upwards, and the slide placed on dry ice for 5 min before removing the coverslip with a razor blade. Slides were air dried at RT, mounted in Hoechst-containing mounting medium, sealed with nail polish, and stored at 4°C before imaging.

### Preparation of *Xenopus* egg extracts and fixed spindle imaging

Species-specific modifications to the *X. laevis* protocol^26,27^ were as follows. *X. tropicalis*: eggs were collected by gentle squeezing in *tropicalis* system water, and EGTA (10 mM) and MgCl_2_ (2 mM) concentrations were doubled for extract preparation. *X. borealis*: eggs were laid overnight in 0.5x MMR, again with EGTA (10 mM) and MgCl_2_ (2 mM) doubled in extract preparation. *X. longipes* eggs were collected by gentle squeezing into *longipes* system water, and extracts prepared in 1.5 mL microcentrifuge tubes using a tabletop centrifuge.

Squash preparations were used for spindle morphometry and bipolarity measurements to preserve morphology. Spindles in metaphase extract (2 µL) was mixed with 3 µL of Hoechst-containing fixative on a precleaned glass slide, gently covered with a coverslip, sealed with nail polish, dried overnight at room termperature in the dark, and stored at 4°C.

For immunofluorescence analysis of spindle proteins, metaphase spindles were processed by spindown as described previously^26^ using rabbit antibodies against TPX2^36^ (1:5000), KIF2A (1:2500, Novus Biologicals), Katanin p60^34^ (1:500), Eg5 (1:2000, Kapoor Lab), XCTK2 (1:1000, Walczak Lab), RCC1^53^ (1:500), and CENPE^54^ (1:100).

Line-scan profiles were analyzed in R (version 2026.07.1) by comparing mean values within each 1% spindle length bin using two-tailed t-tests, with P-values plotted across spindle length to identify significant differences between conditions.

### Live imaging of *Xenopus* egg extract droplets and Ran-GTP gradient visualization

To image spindle assembly and monitor spindle bipolarity, egg extracts were combined with embryo nuclei of varying ploidies, with species-matched controls and mixed-ploidy reactions prepared in parallel. 20 µL extract reactions (200 nuclei/µL) were added to 200 µL of 008-fluorosurfactant (RAN Biotechnologies) and vortexed at max speed for 10 seconds to encapsulate extract in droplets and enrich for diameters in the 60-150 µm range. The resulting droplets were loaded into silane-coated glass capillaries (2000 x 2000 x 150 µm), submerged in ∼2 mL of heavy mineral oil (Sigma-Aldrich) in a well of a glass bottom 1.5 coverslip 6-well plate, and imaged on a ZEISS CD7 microscope Multiple single plane fields were imaged at 60 second intervals for 50 min (*X. tropicalis*) or 60 min (*X. laevis* or *X. longipes*).

For Ran-GTP visualization, the CFP/YFP probe Rango^55^ was added at to extracts at 2 µM before nuclei addition. Reactions were incubated for 45 min (*X. tropicalis*) or 55 min (*X. laevis* and *X. longipes*) and loaded into droplets and glass capillaries as above before acquiring CFP, YFP, and FRET z-stacks (15 µm depth, 1.5 µm steps). FRET was validated by photobleaching. Following background subtraction in FIJI (version 2.16.0), FRET/CFP ratio images were generated and pseudocolored using the gem lookup table. Line scans were measured across 50 x 80 µm regions centered on the metaphase plate, averaged across spindles, and normalized to the corresponding species-matched control.

### Western blotting of spindle proteins

Western blots were performed using three independent extract preparations per *Xenopus* species (nine lanes per protein) with three technical replicates. Extract protein concentration was measured by BCA assay (Thermo Scientific), and samples prepared in 1x Laemmli sample buffer containing 2.5% beta-mercaptoethanol. Proteins (10 µg/lane) were resolved on BioRad Mini-PROTEAN TGX 4-20% gradient gels and transferred to nitrocellulose overnight at 15V in transfer buffer containing 20% methanol. Total transferred protein was quantified using Revert Total Protein Stain on a Li-COR Odyssey CLx. Membranes were blocked in 5% dry milk, incubated with primary antibodies overnight at 4°C, and secondary antibodies (1:10,000) for 2 hours at room temperature. Band intensities were quantified in FIJI. Each lane was normalized to total protein intensity per lane and averaged across biological replicates to determine relative protein abundance among species based on previous mass spectrometry data^56^.

Primary antibodies were rabbit anti-TPX2^35^ (1:5000), mouse anti-Ran (1:2500, BD Biosciences), rabbit anti-KIF2A (1:5000, Novus Biologicals), rabbit anti-Katanin p60^34^ (1;1000), rabbit anti-XMAP215 (1:2500, Mitchison Lab), rabbit anti-Eg5 (1:2000, Kapoor Lab), rabbit anti-importin alpha^57^ (1:1000), rabbit anti-RCC1^53^ (1:2500), rabbit anti-XKCM1(MCAK)^58^ (1:2500) and rabbit anti-KIF18A^44^ (1:2500, Mayer lab)

### Alteration of *X. laevis* egg extracts

To mimic *X. longipes* protein levels, *X. laevis* egg extracts were supplemented with recombinant *X. laevis* TPX2 (60 nM added; ∼200 nM final), and KIF2A (I252S) (30 nM added; ∼85 nM final) in CSF-XB and used for live and fixed imaging. Control extracts received an equivalent volume of CSF-XB.

To activate XKCM1 (MCAK), 200 nM or 500 nM UMK57 (in DMSO) was added to *X. laevis* egg extracts prior to mixing with nuclei. For reactions containing TPX2, these extracts were supplemented with recombinant *X. laevis* TPX2 to ∼200 nM in CSF-XB. Control extracts received an equivalent volume of both DMSO and CSF-XB.

### Transcriptomic analysis of *X. longipes* spindle proteins

Total RNA from an adult *X. longipes* female was isolated through homogenization of multiple tissues (heart, muscle, kidney, skin, liver, and ovary as independent samples) in TRIzol (Thermo Scientific), followed by chloroform addition to isolate nucleic acids. Nucleic acids were bound to a Qiagen RNeasy column through centrifugation, treated with DNase I to selectively degrade DNA, and then eluted and checked for sample purity via Qubit analysis. Total mRNA samples were prepped for sequencing using PacBio Kinnex array methodology and run on a PacBio Revio 25M SMRTcell. Resulting long-read data was used to generate a *de novo* transcriptome assembly, using *X. laevis* and *X. tropicalis* sequences as reference (given no genomic data is currently available for *X. longipes*). Individual *X. longipes* protein sequences shown in Figure 4F were aligned against other *Xenopus* species using ClustalOmega.

## Acknowledgments

We would like to thank the numerous laboratories and scientists who have contributed reagents and developed techniques used in this project. We thank Kelly Miller, Clotilde Cadart, Coral Zhou, and Ziyuan Liu for their contributions to establishing *X. longipes* as a laboratory research model. We also acknowledge Amarisa Gonzalez and Aidan Heller for caring for *X. longipes* tadpoles and contributing to data acquisition and analysis of spindle morphometrics. The Rango FRET probe was developed by Petr Kalab, the lipid droplet approach was adapted from methods developed by the Gelens Lab (KU Leuven), and antibodies against numerous proteins were generously provided by members of the *Xenopus* community, including the Mitchison, Kapoor, Mayer, and Walczak Labs. We thank past and present Heald Lab members for valuable discussions and insights throughout this project. We also thank Anuradha Mukherjee, Alex Lessenger, Megan Onyundo, and Helena Cantwell for helpful feedback on the manuscript. No AGI or LLM was implemented in the process of acquiring and analyzing the data of this project.

## Funding

National Institutes of Health MIRA grant R35GM118183 (RH)

Flora Lamson Hewlett Chair in Biochemistry (RH)

Damon Runyon Cancer Research Foundation – Merck Fellow DRG 2503-23 (GC)

Belgian American Education Foundation Research Fellowship (LP)

National Research Foundation of Korea RS-2023-NR076754 (TK)

## Author contributions

Conceptualization: GC, RH

Methodology: GC, CC, LP, TK, RH

Investigation: GC, CC, LP, DW

Visualization: GC, CC, LP, DW

Funding acquisition: GC, LP, RH, TK

Project administration: GC, RH

Supervision: TK, RH

Writing – original draft: GC, RH

Writing – review & editing: GC, CC, LP, DW, TK, RH

---

**Fig S1.**
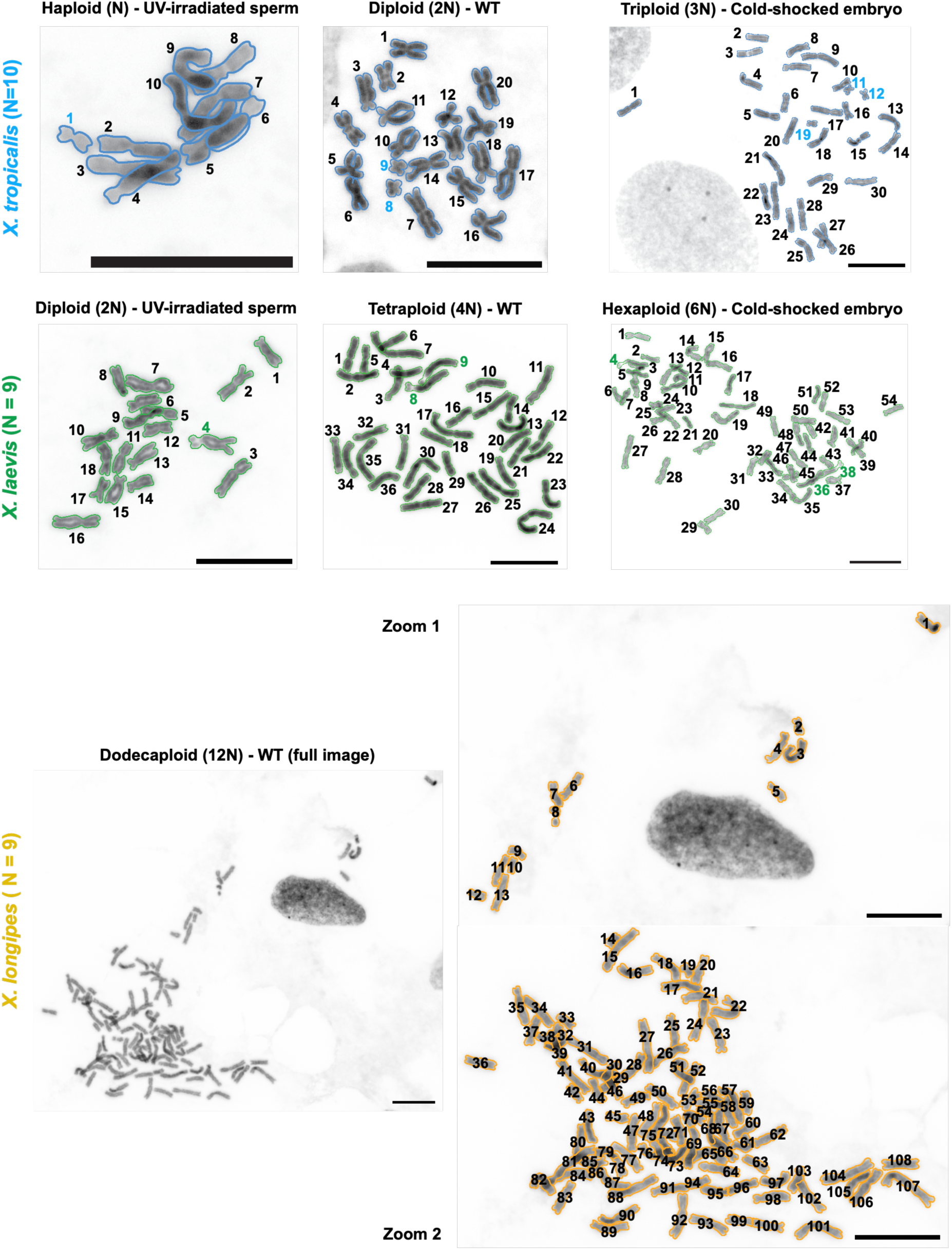
Validation of embryo nuclear ploidy by metaphase spreads. Metaphase spreads of tailbud cells of *Xenopus* tadpoles (NF stages 35-40) derived from the same clutches used for isolation of embryo nuclei. Each chromosome is outlined and numbered to aid in visualization (numbers do not correspond to chromosome identity). Colored labels mark cytologically distinctive reference chromosomes used to confirm ploidy in addition to total chromosome number (chromosome 10 in *X. tropicalis* and chromosome 3L in *X. laevis*). Scale bar, 10 µm. Because a chromosome fusion proceeded the hybridization and whole-genome duplication event that gave rise to *X. laevis*, haploid chromosome numbers are 10 for *X. tropicalis* and 18 for *X. laevis*^52^. Haploid embryos were generated using UV-irradiating sperm for fertilization, and triploid embryos by cold-shock after fertilization to prevent second polar body extrusion.Supplemental Figure 2

**Fig S2.**
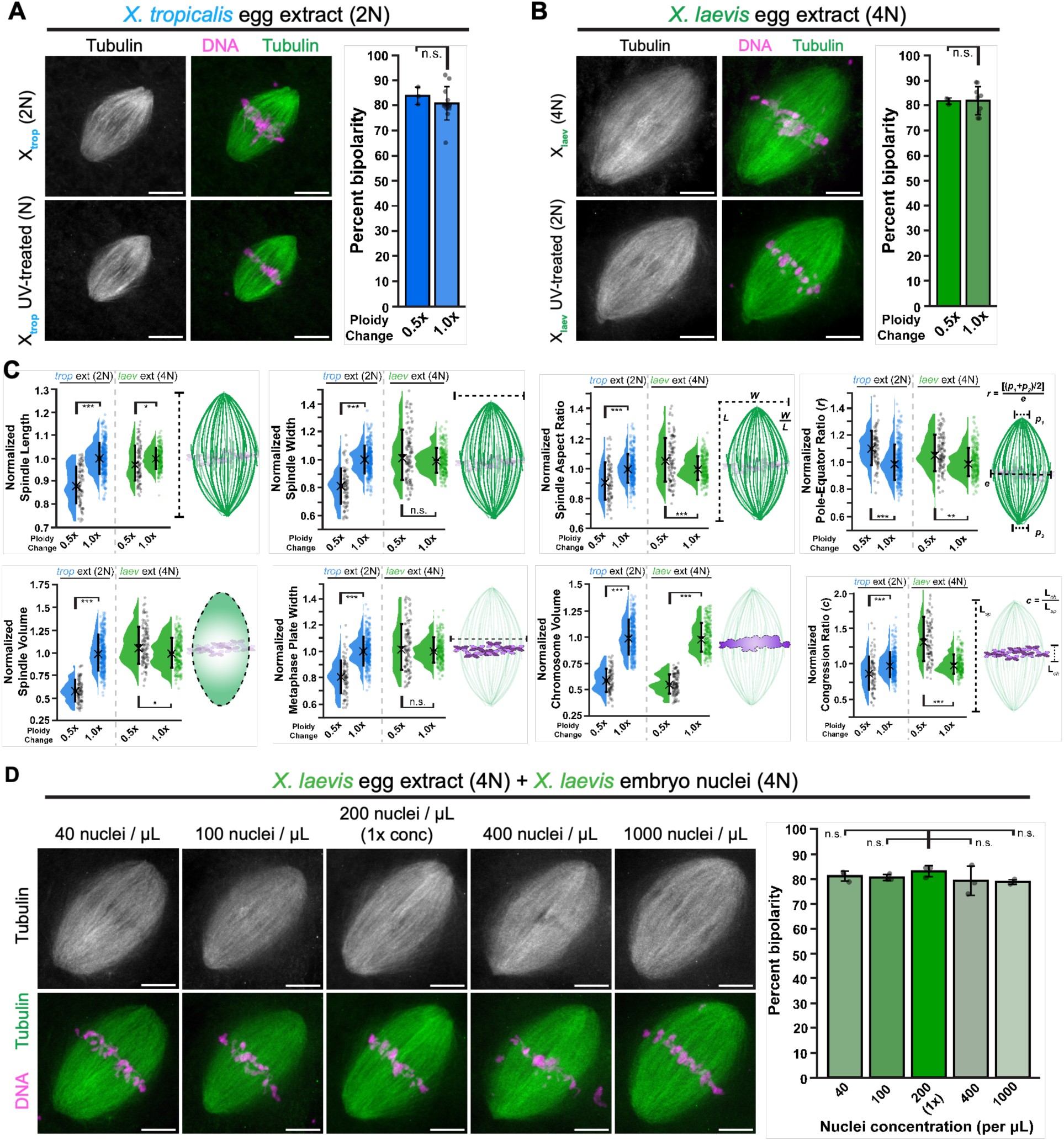
Spindle bipolarity is unaffected by reduced ploidy or altered nuclear concentration. **(A,B)** Fixed spindles assembled in *X. tropicalis* and *X. laevis* egg extracts formed with control or half-ploidy nuclei. Individual points represent biological replicates (≥3 per condition and ≥30 spindles per replicate. Bars indicate the mean ± standard deviation. **(C)** Quantification of spindle morphometrics observed in (A,B). Each point represents a single spindle. Bars indicate the mean ± standard deviation, and crosses indicate the median. Asterisks denote p values determined by ANOVA; n.s., not significant. Spindle numbers for each measurement are provided in Table S1. **(D)** Fixed spindle squashes from X. *laevis* extract combined with nuclei of the same ploidy at varying concentrations (200 nuclei/μL was used as the standard concentration throughout this study). Individual points represent biological replicates (≥3 per condition and ≥20 spindles per replicate). Bars indicate the mean ± standard deviation. All images are maximum projections of z-stacks. Scale bar, 10 µm. Significance was determined by ANOVA. The number of spindles in each condition is provided in Table S1.

**Fig S3.**
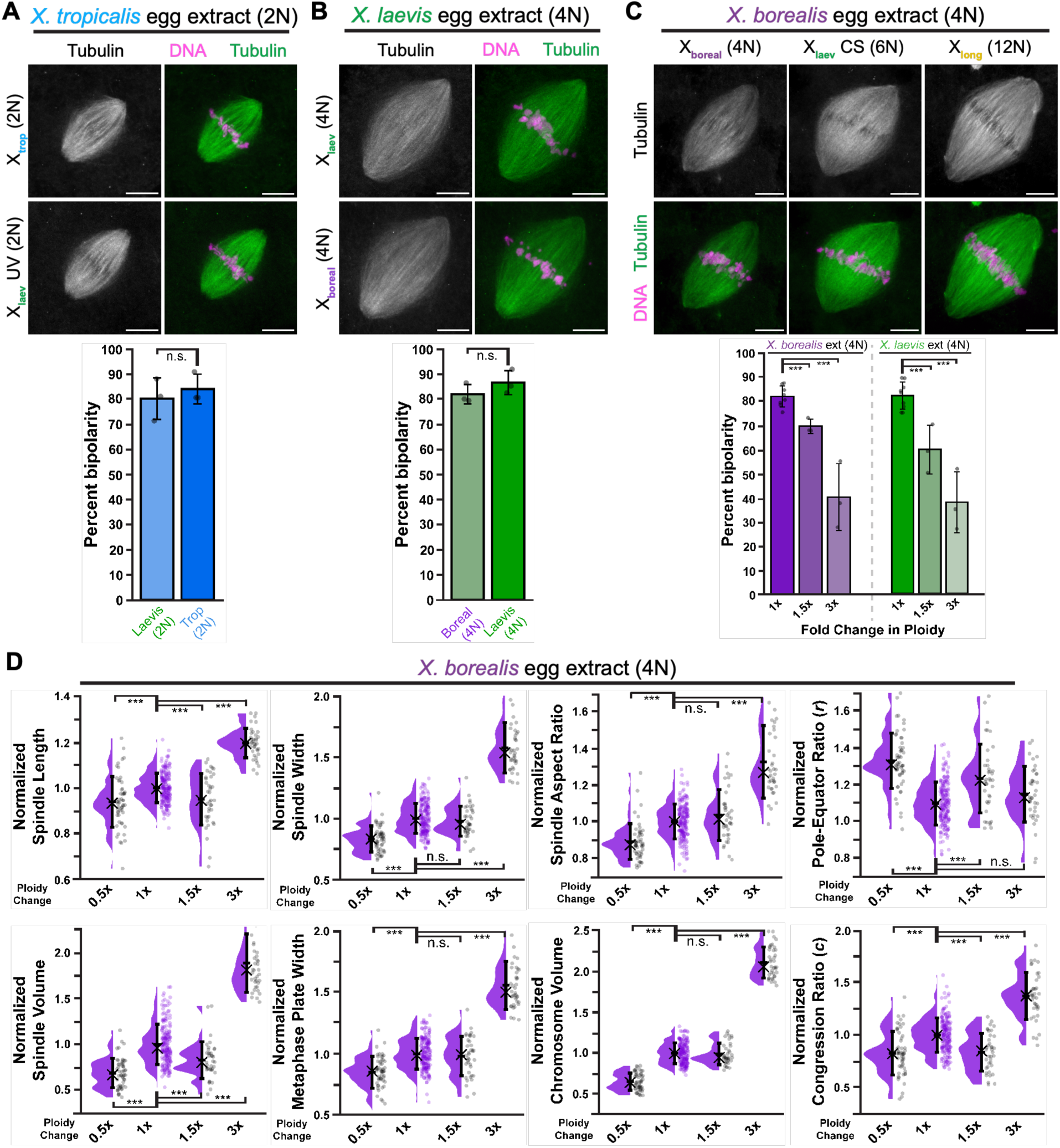
Ploidy, rather than species origin, affects spindle bipolarity. **(A)** Fixed spindle squashes and percent bipolarity of *X. tropicalis* egg extracts combined with 2N *X. tropicalis* or haploid-derived 2N *X. laevis* nuclei. **(B)** Fixed spindle squashes and percent bipolarity of *X. laevis* egg extracts combined with 4N *X. laevis* or 4N *X. borealis* nuclei. **(C)** Fixed spindle squashes and percent bipolarity of *X. borealis* extracts combined with nuclei from 4N *X. borealis*, triploid-derived 6N *X. laevis*, or 12N *X. longipes*. Percent bipolarity measurements for *X. laevis* were adapted from Figure 1B for direct comparison between the two tetraploid *Xenopus* species. **(D)** All eight spindle morphometrics measured from *X. borealis* mixed-ploidy spindles shown in (C). Each point represents a single spindle. Data are pooled from at least three biological replicates (≥40 spindles per replicate). Bars indicate the mean ± standard deviation, and crosses indicate the median. Asterisks denote p values determined by ANOVA; n.s., not significant. Spindle numbers for each measurement are provided in Table S1. All images are maximum projections of z-stacks. Scale bar, 10 µm. Individual points in all bipolarity graphs represent biological replicates (≥3 per condition and ≥20 spindles per replicate). Bars indicate the mean ± standard deviation. Significance was determined by ANOVA.

**Fig S4.**
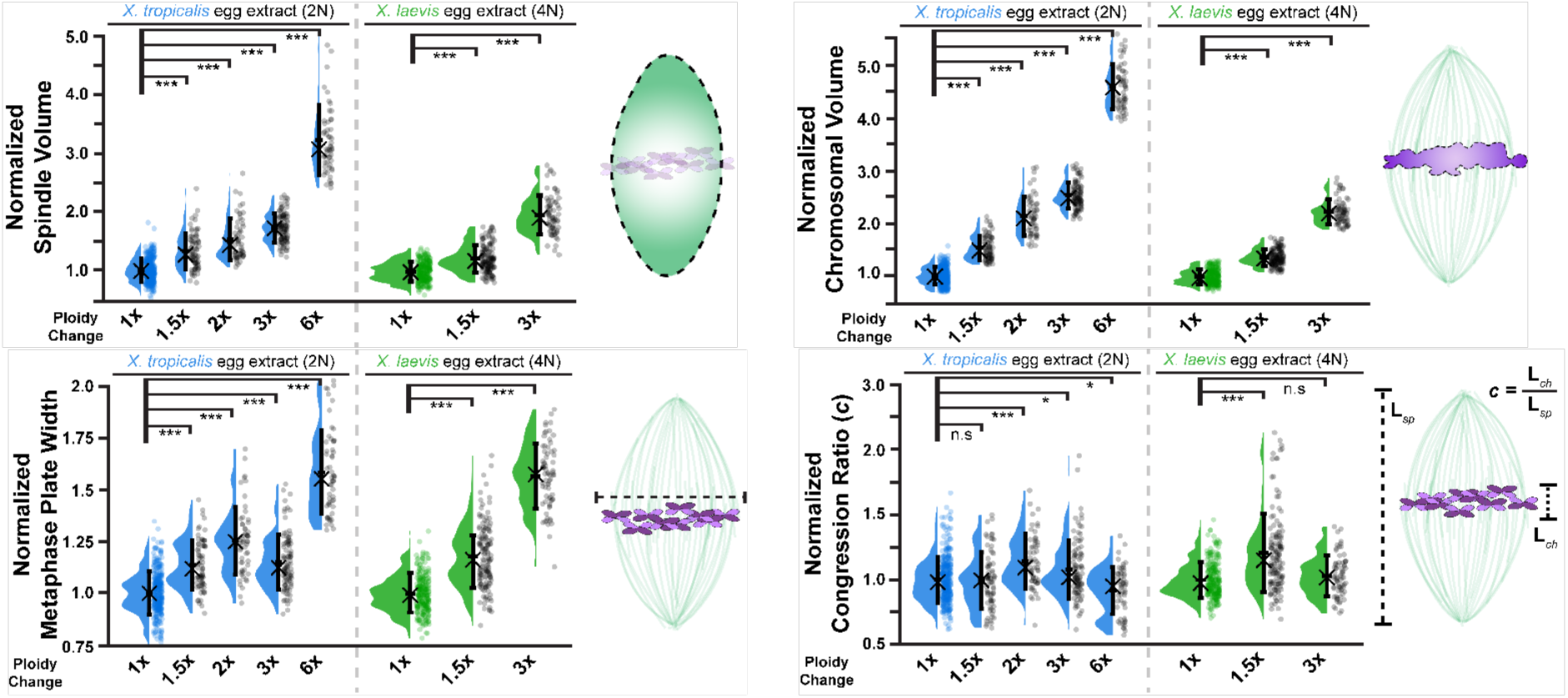
Additional spindle morphometrics measured in mixed-ploidy spindles. Four additional measurements of bipolar spindle morphology, including spindle volume, metaphase plate width, metaphase plate volume, and the fraction of pole-to-pole distance occupied by chromosomes, as a measure chromosome congression. Each point represents a single spindle. Data are pooled from at least three biological replicates (≥55 spindles per replicate). Bars indicate the mean ± standard deviation, and crosses indicate the median. Asterisks denote p values determined by ANOVA; n.s., not significant. The number of spindles analyzed for each measurement is provided in Table S1.

**Fig S5.**
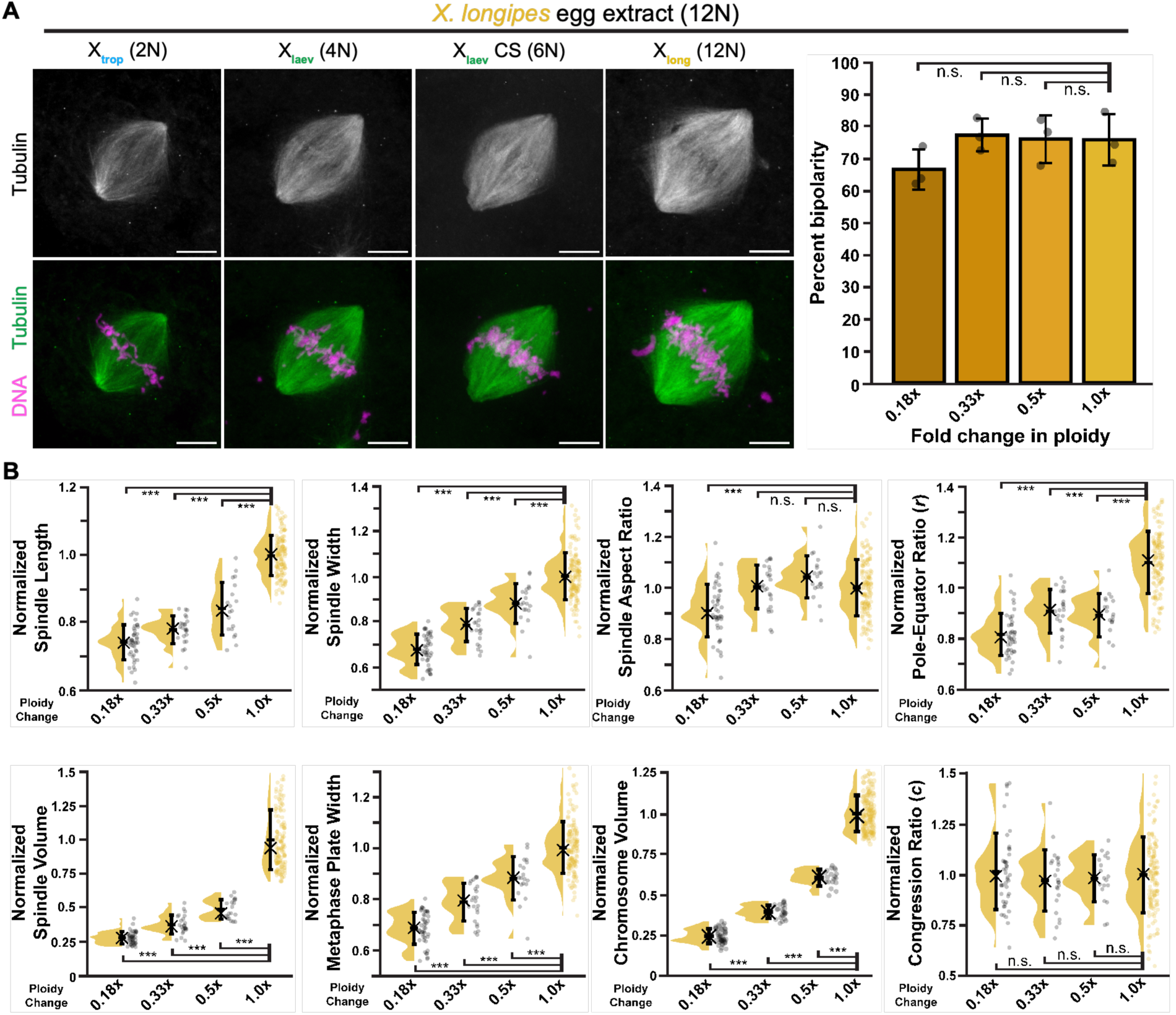
Reductions in ploidy do not significantly impact the efficiency of bipolar spindle assembly. **(A)** Fixed squashes of *X. longipes* extract spindles assembled around nuclei of decreasing ploidy. All images are maximum projections of z-stacks. Scale bar, 10 µm. Shaded points show individual replicates (≥35 spindles per replicate); bars indicate mean value ± standard deviation. **(B)** All eight spindle morphometrics measured in the bipolar spindles shown in (A). Points represent individual spindles, pooled across three biological replicates (≥20 spindles per replicate). Bars indicate the mean ± standard deviation, and crosses indicate the median. Asterisks denote p values determined by ANOVA. The number of spindles measured is provided in Table S1.

**Fig S6.**
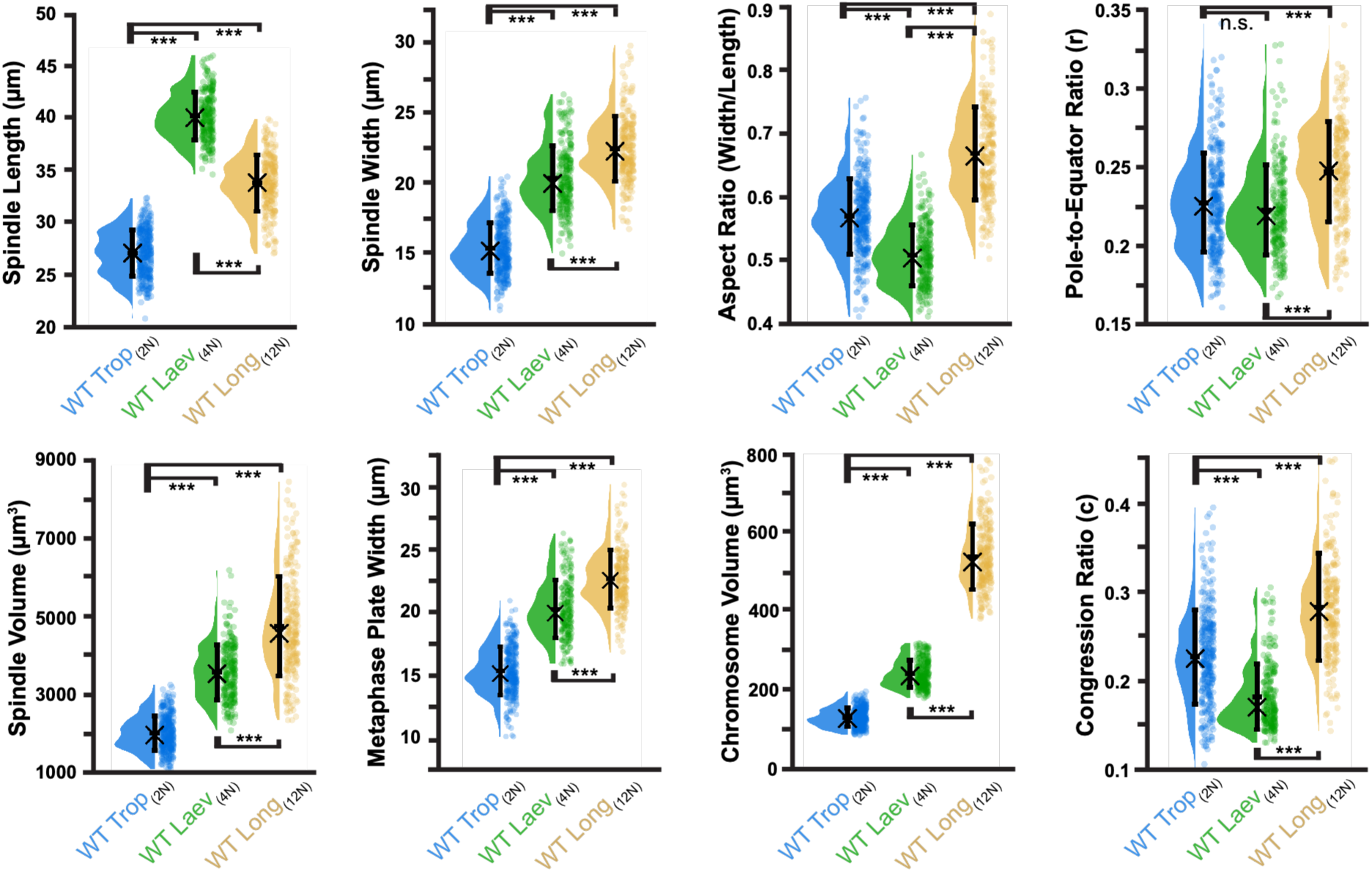
Comparison of wild-type *Xenopus* spindle morphometrics across *X. tropicalis, X. laevis,* and *X. longipes*. Comparison of all eight native wild-type egg extract spindle morphometrics across the three *Xenopus* species. Individual points represent single spindles, pooled from at least three biological replicates (≥20 spindles per replicate). Bar indicates the mean value ± standard deviation, and crosses indicate the median. Asterisks denote p values determined by ANOVA. The number of spindles measured is provided in Table S1.

**Fig S7.**
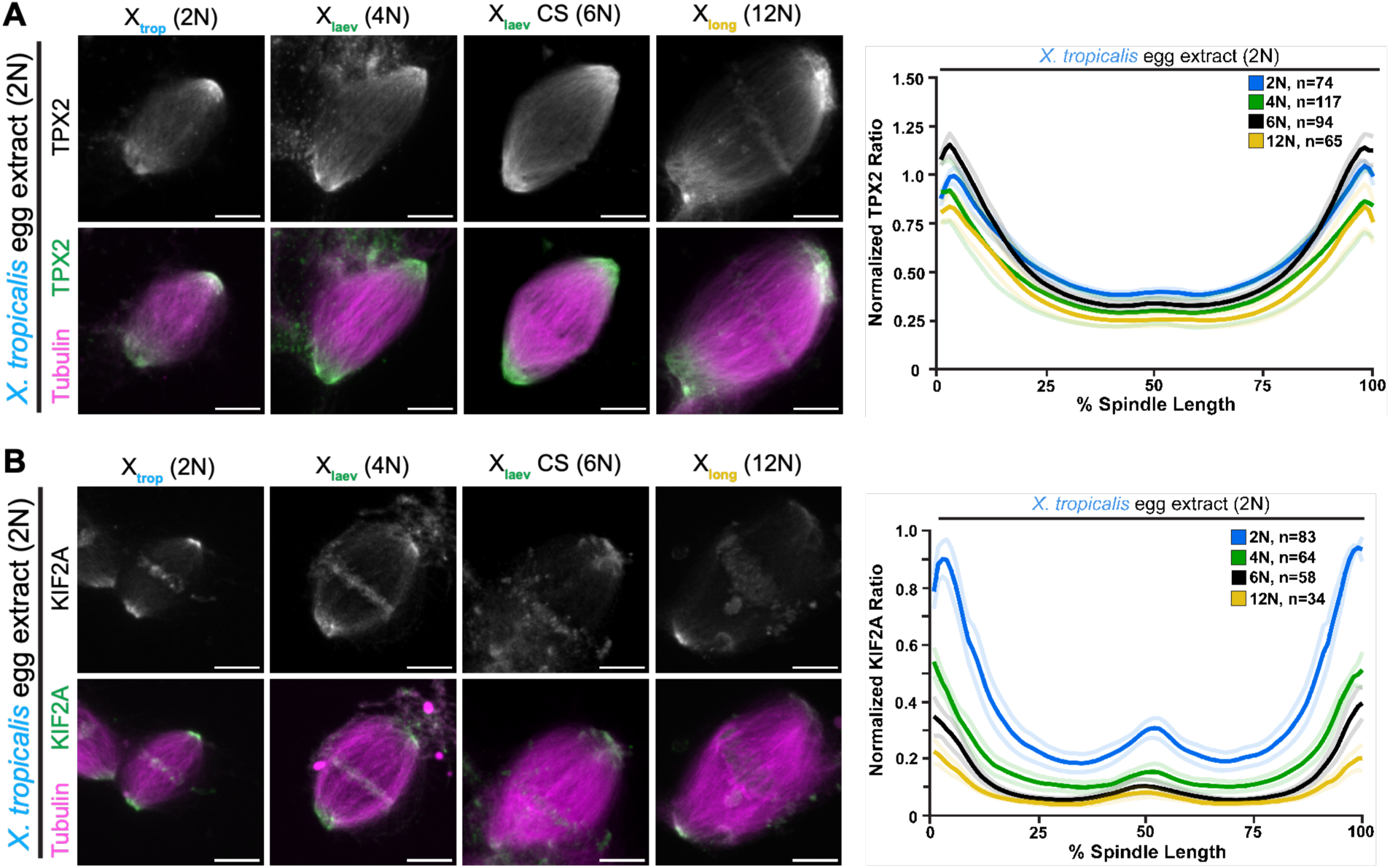
Linescan analysis of TPX2 and KIF2A in *X. tropicalis* mixed-ploidy spindles. **(A,B)** Linescan analysis of TPX2 and KIF2A intensity relative to spindle microtubule intensity along the pole-to-pole axis in *X. tropicalis* extracts, normalized to species-matched control. Solid lines represent the average profile, shaded lines of the same color represent ± SEM. The number of individual linescans in each condition is shown. Representative images for all panels are maximum projections of z-stacks. Scale bar, 10 µm.

**Fig S8.**
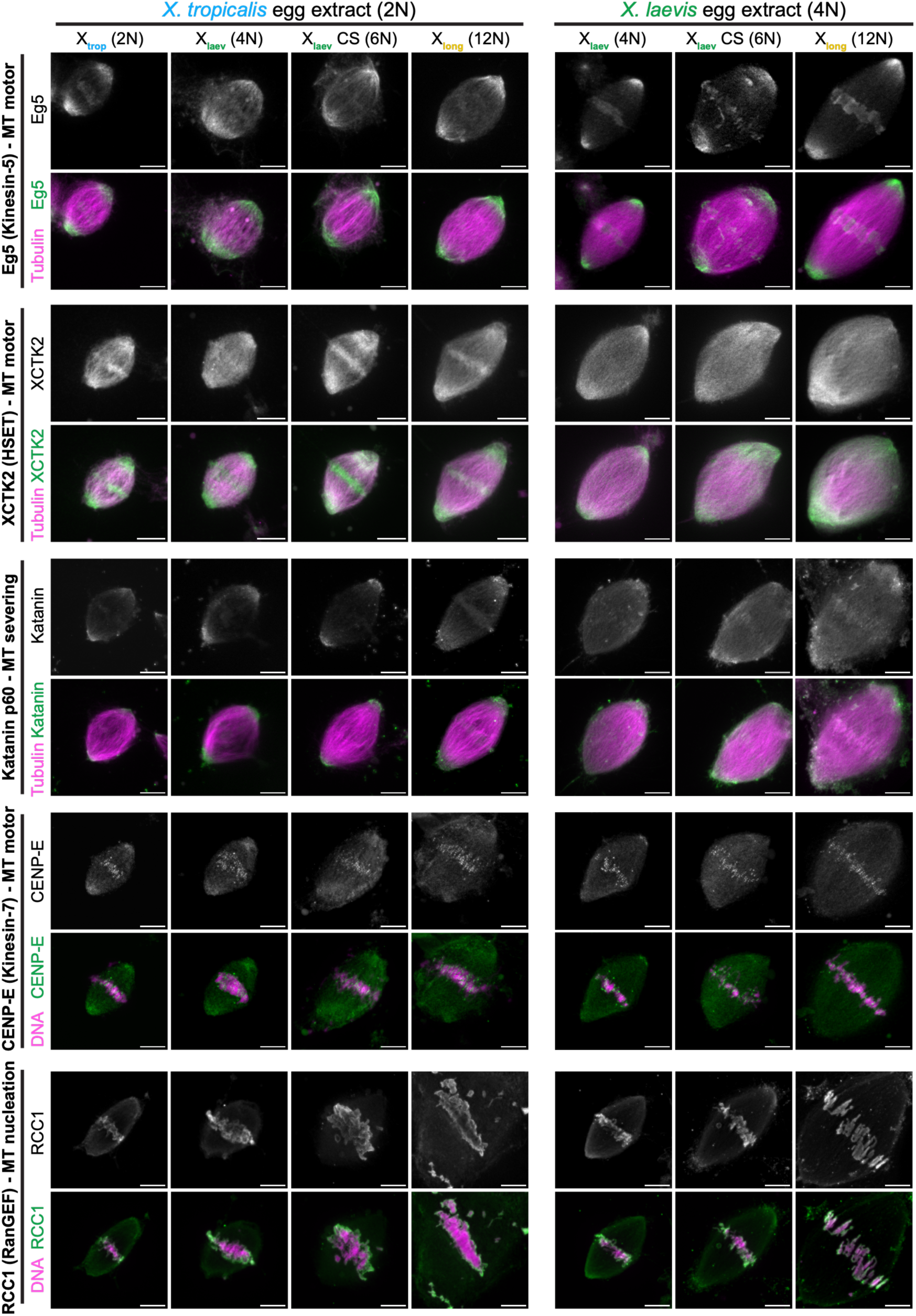
Localization of five additional spindle-associated proteins with increased ploidy. Immunofluorescence imaging of five additional proteins in mixed-ploidy spindles in *X. tropicalis* and *X. laevis* extracts. A classification of each protein’s role in spindle assembly is provided. Representative images for all conditions are maximum projections of z-stacks. Scale bar, 10 µm.

**Fig S9.**
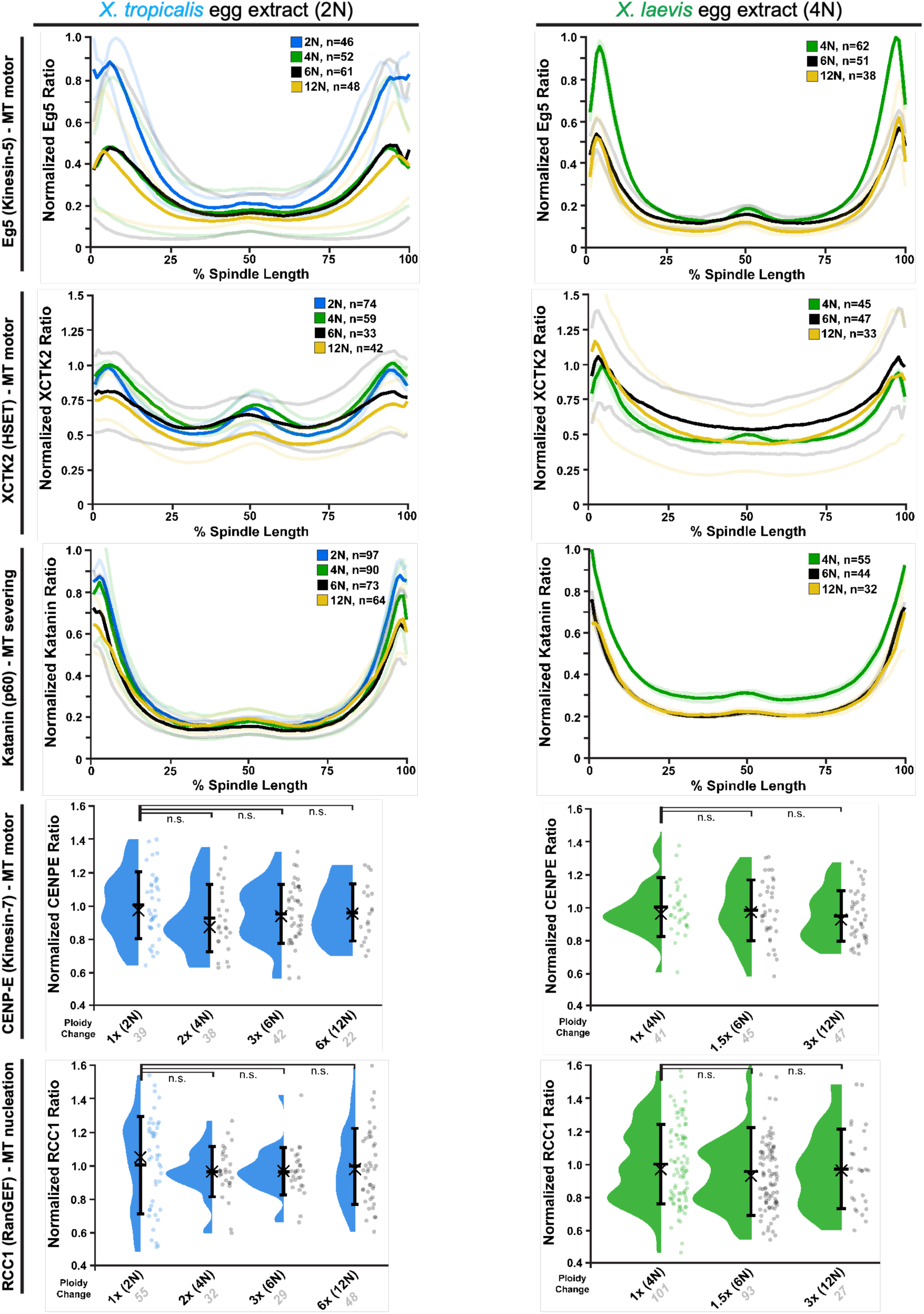
Linescan and ROI analysis of five additional spindle proteins with increased ploidy. Linescan (Eg5, XCTK2, and Katanin p60) or ROI (RCC1 and CENPE) analyses of the five proteins shown in Figure S8. Linescan data were normalized to the tubulin intensity across the spindle, while ROI data were normalized to total DNA intensity at the metaphase plate. For linescan analysis, solid lines represent the average profile, and shaded lines represent ± SEM. The total number of spindles in each condition is shown. For ROI analysis, each point represents a single spindle. Data are pooled from at least three biological replicates (≥10 spindles per replicate). Bars indicate the mean ± standard deviation, and crosses indicate the median. Asterisks denote p values determined by ANOVA; n.s., not significant. The total number of images in each condition is shown on the x-axis.

**Fig S10.**
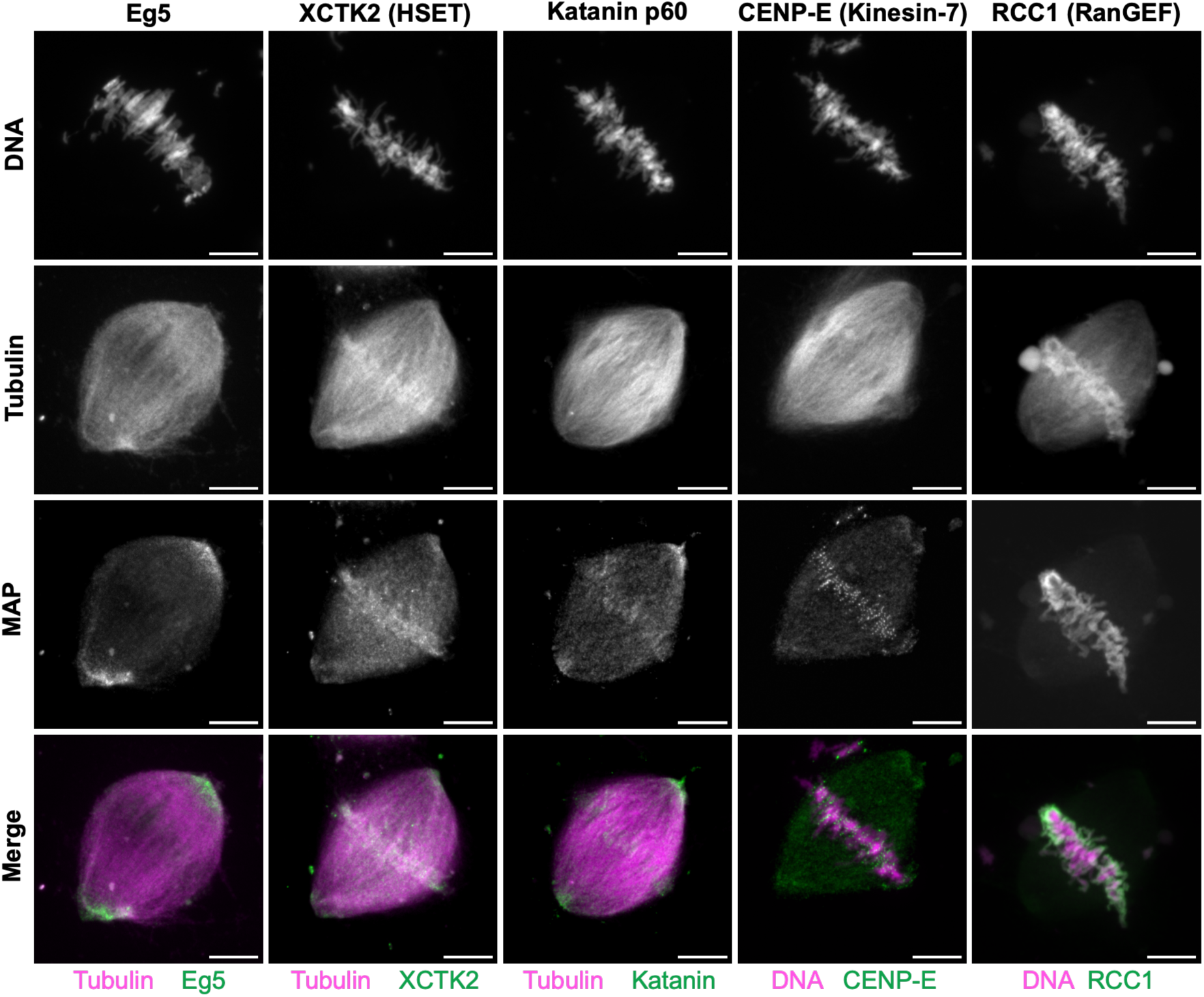
Localization of various spindle-associated proteins in native *X. longipes* spindles. Representative images showing the localization of proteins within native *X. longipes* extract spindles. All images are maximum projections of z-stacks. Scale bar, 10 µm.

**Fig S11.**
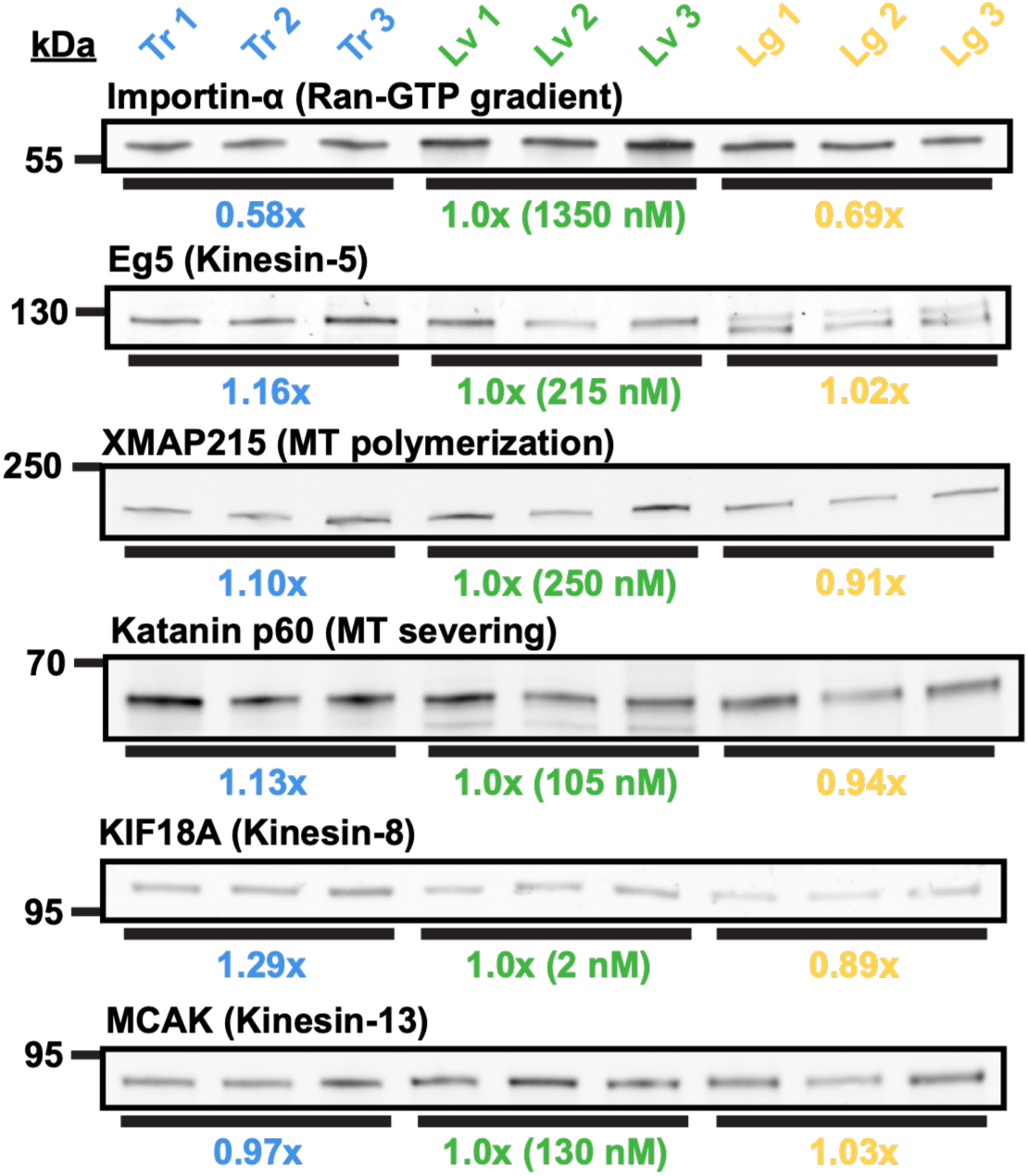
Many spindle-associated proteins have similar levels across *Xenopus* species. Western blots showing the levels of six additional spindle-associated proteins across *Xenopus* egg extracts. Each lane contained 10 μg total protein and represents a biological replicate, and all blots were repeated three times. Band intensities were normalized to the mean intensity in *X. laevis* extracts; estimated protein concentrations are shown in parentheses.

**Table S1. Individual vales for all morphometrics and percent bipolarity measured across all combinations of nuclei and extract** Mean values and standard deviations measured for all eight spindle morphometrics and percent bipolarity measurements across all combinations of extracts and nuclei presented in this study. The total number of spindles in each condition is also included.

